# Lepbase: the Lepidopteran genome database

**DOI:** 10.1101/056994

**Authors:** Richard J Challi, Sujai Kumar, Kanchon K Dasmahapatra, Chris D Jiggins, Mark Blaxter

## Abstract

As the generation and use of genomic datasets is becoming increasingly common in all areas of biology, the need for resources to collate, analyse and present data from independent (Tier 1) species-level genome projects into well supported clade-oriented (Tier 2) databases and provide a mechanism for these data to be propagated to pan-taxonomic (Tier 3) databases is becoming more pressing. Lepbase is a Tier 2 genomic resource for the Lepidoptera, supporting a research community using genomic approaches to understand evolution, speciation, olfaction, behaviour and pesticide resistance in a wide range of target species. Lepbase offers a core set of tools to make genomic data widely accessible including an Ensembl genome browser, text and sequence homology searches and bulk downloads of consistently presented and formatted datasets. As a part of the taxonomic community that we serve, we are working directly with Lepidoptera researchers to prioritise analyses and add tools that will be of most value to current research questions.

## Findings

### Background

With the falling cost of data generation and the corresponding increase in the rate of genome sequencing (Muir et al. 2016), use of large-scale genomic datasets is increasingly common (e.g. (Brawand et al. 2014; Soria-Carrasco et al. 2014; Zhang et al. 2014; Foote et al. 2015)). Genome sequencing projects are no longer the preserve of dedicated sequencing centers or large scale collaborations but are increasingly instigated and completed by single laboratories, or small groups of collaborators (e.g. (Cong et al. 2016)). The goals of such sequencing projects have also diverged and, at the extremes, some projects strive for reference quality, chromosomal assembly (Vij et al. 2016), while others use genomics as a means to target specific genes or gene families.

The importance of making genomic data publicly available to support further research on the same or related species has been well established (Toronto International Data Release Workshop Authors et al. 2009). It is common practice for labs that generate genomic datasets to deposit the raw data to a public archive such as GenBank, ENA, or DDBJ, and either to deposit the assemblies and annotations alongside the raw data or to host them along with additional analysis files on a dedicated server. While these steps make the data available, they fall short of making the data widely accessible. Genome databases built by individual laboratories for a single focal species can be brittle, hard to update, and difficult to maintain in the face of short term funding cycles. Considerable bioinformatics input is required to make use of these resources (Muir et al. 2016), and particularly in comparative studies where data from many individual projects may be combined.

For well studied organisms, mature portals exist that provide rich contextual information for genes and other genomic features (e.g. (Rhee 2003; Bult et al. 2015; Attrill et al. 2016; Howe et al. 2016)). Most projects lack the resources and/or expertise to develop, host and maintain these high-performance tools. Conversely, large, pan-taxonomic aggregative databases, such as Ensembl (Yates et al. 2016), while hosting large-scale comparative resources (Herrero et al. 2016), lack the domain-specific knowledge to meet the specific requirements of taxon-oriented communities and are often unable to incorporate data that do not meet specific quality criteria for inclusion.

An important model for the growth of genomic research in non-model organisms is to establish community-based, taxon-oriented genomic databases and resources as the middle of three tiers, aggregating, analysing and displaying data from lab-scale (Tier 1) projects and ultimately providing a conduit for these data to reach the large, pan-taxonomic (Tier 3) databases (Parkhill et al. 2010). While the largest taxon-oriented databases (such as FlyBase (Attrill et al. 2016) or WormBase (Howe et al. 2016)) gain stability through core funding due to the importance of these species as model organisms, here we consider Tier 2 databases to be those that achieve stability through offering a defined mechanism for propagation to pan-taxonomic Tier 3 databases through use of a compatible architecture. This definition excludes some taxon-oriented resources that do not fall within the three-tiered hierarchy, such as those based on the GMOD infrastructure (Generic Model Organism Database (GMOD) http://gmod.org/), which is open source and community-driven but best suited to use in Tier 1 databases. Existing Tier 2 databases for non-models (e.g. WormBase Parasite (Howe et al. 2016), VectorBase (Giraldo-Calderon et al. 2015), (PomBase McDowall et al. 2015) and Avianbase (Eöry et al. 2015)) are typically based on the Ensembl (Yates et al. 2016) architecture as this is currently the most complete centrally supported genome-centered databasing/viewing platform.

The lepidopteran research community is making growing use of genomic data in understanding evolution (Heliconius Genome Consortium 2012; Ahola et al. 2014; Derks et al. 2015), speciation (Cong et al. 2015b; Cong et al. 2016), olfaction (You et al. 2013; Tanaka et al. 2009; Zhan et al. 2011), behaviour (Derks et al. 2015; You et al. 2013) and pesticide resistance (You et al. 2013) in a wide range of target species. We set out to build a genomics database for this community that would be rapidly responsive to new data, particularly from new species. We aimed to include not just a single reference for each species, but maintain access to older versions and include alternate versions derived from different wild or captive populations. We wanted to be able to include genome assemblies not usually considered for inclusion in aggregative databases because of low quality or the small numbers of people specifically interested in that taxon. We also wanted to normalise annotation across genomes so that comparative genomics would discover biological insight rather than expose methodological differences.

Lepbase is a Tier 2 genomic resource for the Lepidoptera, developed from within the Lepidoptera research community. Here we present the current status of Lepbase, introduce some of the design decisions we have made to facilitate data accession and data access, and outline our plans for the future. We present Lepbase as an example of the suite of tools and services needed to provide a Tier 2 resource to a community.

### Tools & Resources

The core Lepbase toolset includes an Ensembl (Yates et al. 2016) instance, a SequenceServer (Priyam et al. 2015) powered BLAST (Altschul 2014) server, a Web Apollo (Lee et al. 2013) instance and a download server. This allows us to deliver many of the general use cases for taxon-oriented data exploration and analysis. In the development of Lepbase to date, we have made choices and written code to facilitate the creation and sharing of new resources and the addition of new data to existing tools to make Lepbase as responsive as possible to the requirements of the community. As Lepbase matures we are working closely with the Lepidoptera research community to add additional tools, resources and analyses to support more specific questions in Lepidoptera research.

#### lepbase.org

Individual tools and resources may be accessed directly or through our main portal at http://lepbase.org. This uses a customised WordPress (http://wordpress.com) theme to allow flexible categorisation and linking of related tools, analyses and downloads. Announcement of new resources simply requires writing a new post, speeding the process of deployment. Coupled with announcements of major features on existing Lepidopteran and arthropod genomic mailing lists and further announcements from the @lepbase twitter handle, this improves our engagement with the wider community.

#### ensembl.lepbase.org

In line with other Tier 2 databases, we use an Ensembl genomic database and web server (Yates et al. 2016). In the current release (v2, February 2016), our Ensembl database contains 21 assemblies across 17 species (Table 1) including four assemblies imported directly from Ensembl Metazoa (Kersey et al. 2016) *(Bombyx mori (The International Silkworm Genome & The International Silkworm Genome 2008)*, *Danaus plexippus (Zhan et al. 2011)*, *Heliconius melpomene* (Heliconius Genome Consortium 2012) and *Melitaea cinxia* (Ahola et al. 2014)). For *D. plexippus* (Zhan et al. 2014) and *H. melpomene* (Davey et al. 2016) we also host recent improved assemblies.

**Table 1.**
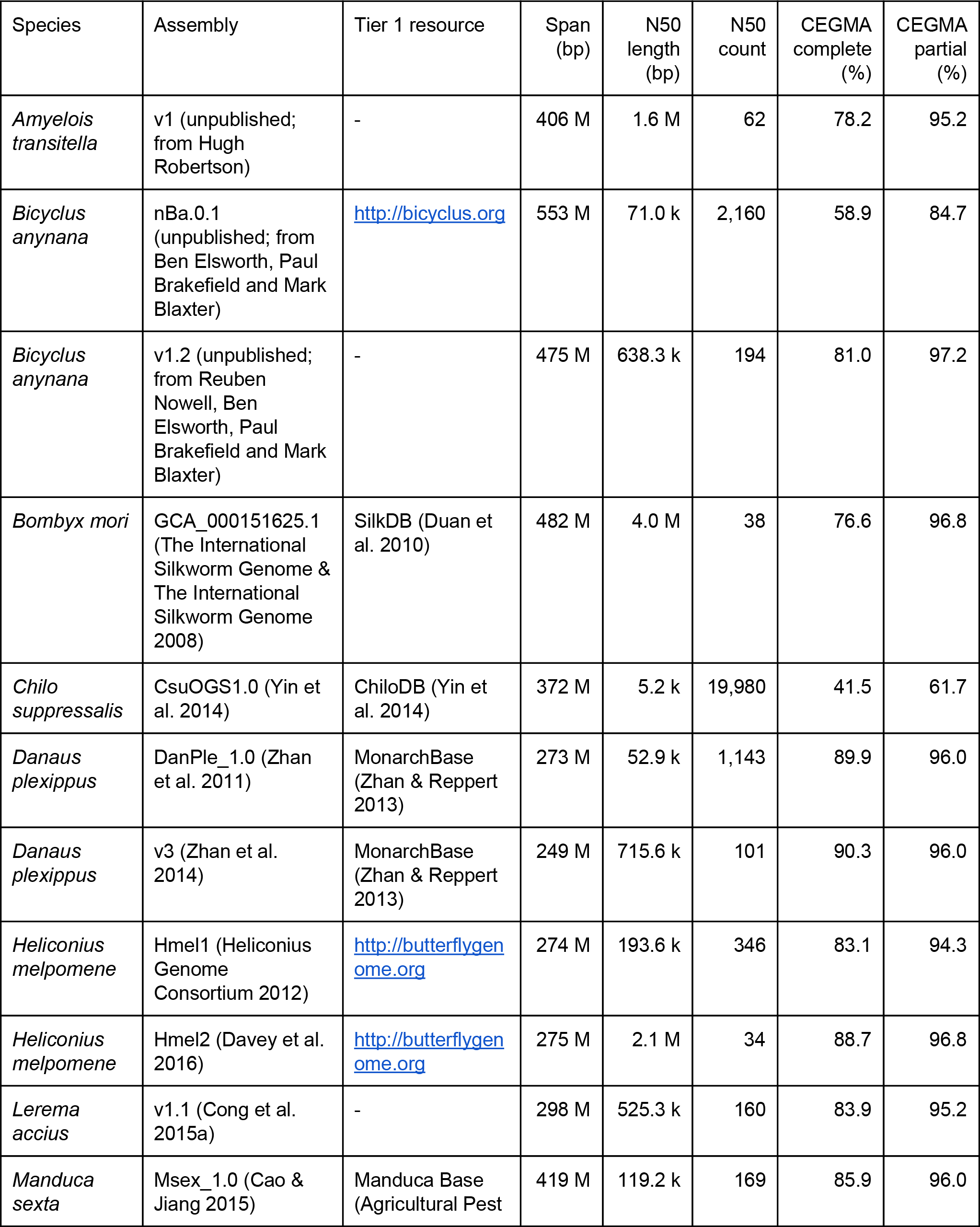
Lepidoptera species and assembly versions available in Lepbase (release v2).

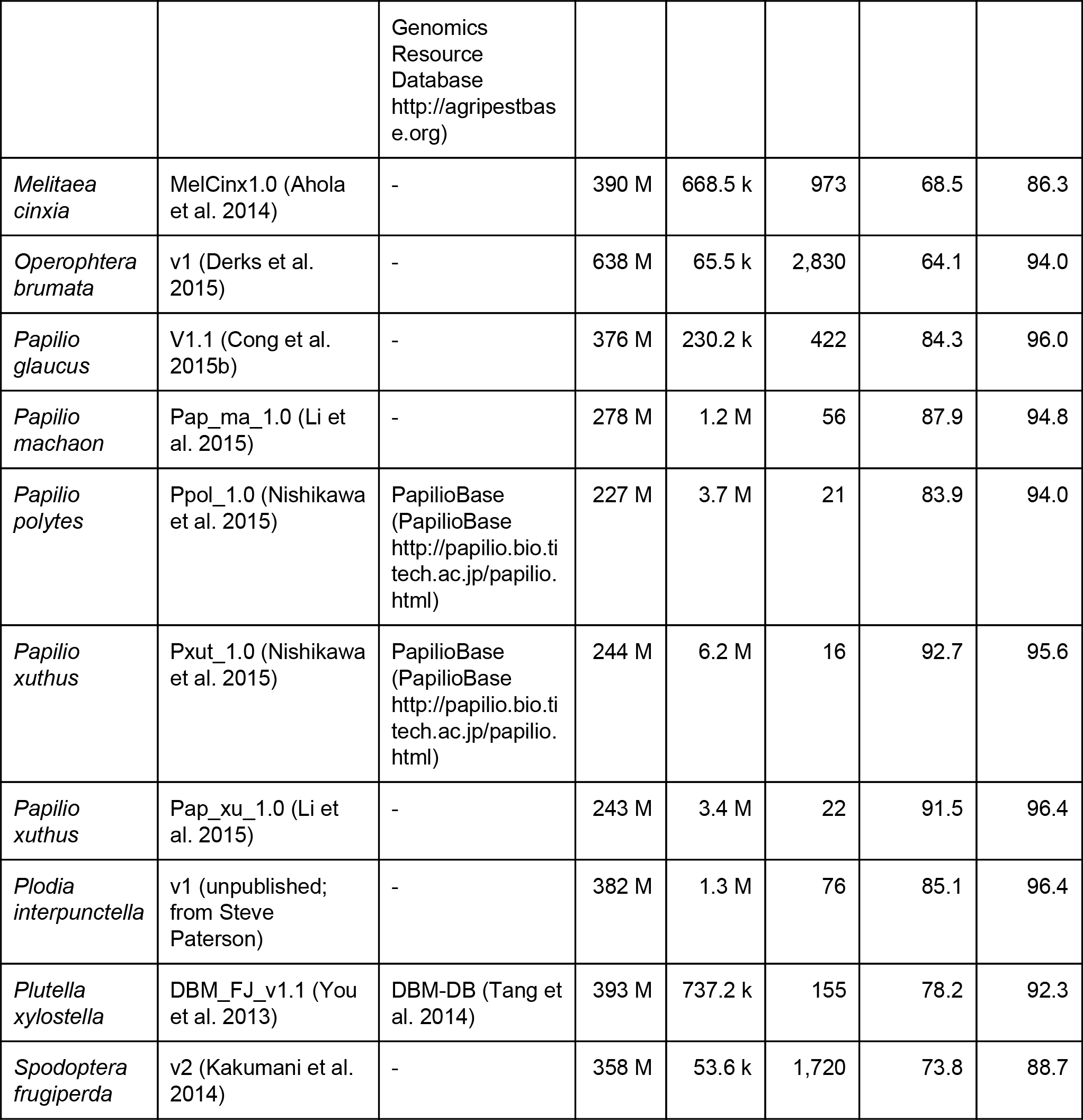

For each species, the Ensembl web interface allows users to view genes and other sequence regions in a genomic context with a rich set of annotations provided by standardised analyses that we apply to each genome (see Analyses). We have also exposed the Ensembl API to allow programmatic access to all of the data for these species. This will allow us to develop novel views of the data and for users to obtain custom datasets through scripted analyses. Use of the API requires a certain level of bioinformatics knowledge so we encourage users with specific requirements to contact us to help develop custom analyses.

We also host sequence data for 18 additional heliconiines (Nymphalidae) sequenced and assembled using the DISCOVAR protocol (Weisenfeld et al. 2014) which have been treated separately due to overall lower quality of the assemblies. This highlights one of the benefits of a community-centered resource such as Lepbase: we are able to accommodate data of differing qualities and maximise the benefits of accessibility - especially for data that would be unlikely to meet criteria for inclusion in pan-taxonomic databases.

We have developed solutions to the challenges presented by deploying an Ensembl instance outside the EBI for a large set of species with heterogeneous data (Challis et al. in prep) which should prove valuable to others wishing to replicate the Lepbase model for other taxa. The general customisations to ensembl.lepbase.org, the search functionality and linkout to blast.lepbase.org have each been implemented as open source Ensembl plugins to ensure that individual components can be reused by other projects.

#### blast.lepbase.org

Identification of potential homologues though sequence similarity search is one of the most important entry points to a genomic database. While Ensembl has the option for built in BLAST functionality, we have chosen to implement an external BLAST server using SequenceServer (Priyam et al. 2015) as we believe that the improved interface best serves the needs of our users. Through our Ensembl BLAST plugin and open source modifications to SequenceServer to allow POST data, we are able to maintain a roundtrip from discovery of a gene in ensembl.lepbase.org through to BLAST searching against the sequence datasets in Lepbase and returning to view results in a genomic context. SequenceServer simplifies the process of adding sequence datasets to a web-accessible BLAST server and we have written open source customisations to allow hierarchical taxon display in SequenceServer that make it straightforward to perform BLAST searches against combined databases from any set of assemblies. We host BLAST databases of scaffold sequences for each of the species on ensembl.lepbase.org and, where available, cDNA and protein databases as well.

#### webapollo.lepbase.org

Web Apollo (Giraldo-Calderon et al. 2015) is a jBrowse (Skinner et al. 2009) based system that facilitates distributed manual editing of gene models on an assembly based on a variety of evidence tracks. Uptake of Web Apollo requires enthusiasm and commitment from a group of potential annotators so we only offer a Web Apollo instance for an assembly when requested by the community. At present we host data for *Heliconius melpomene* assembly Hmel2 (Davey et al. 2016) and for a *Heliconius erato* assembly (Van Belleghem et al. in prep) as part of ongoing (re)annotation efforts in these species.

#### download.lepbase.org

One of the most basic goals for a taxon-oriented resource is to provide access to consistently named and formatted data files. A home page on download.lepbase.org provides links to the major data categories available but beyond that, ease of maintenance is ensured by using automatic directory listings to show which specific data are available. We make relevant raw data and derived analysis files (such as sequence similarity and domain similarities) available for direct download. We use our Ensembl import pipeline (Challis et al. in prep) to verify and repair inconsistencies in provided assembly and annotation files, and make these standardised files (exported from our Ensembl instance) available for download. This assists users in obtaining bulk datasets for analysis and facilitates large-scale comparative analysis in particular.

#### Analyses

Differences between taxa in assembly statistics (e.g. percent repeat content) can arise through differences in program and parameter selection. As an aggregative genomic resource, the standardised datasets at Lepbase provide an opportunity to undertake analyses across all assemblies according to standardised protocols and to make these results available to the wider community. We currently decorate the genomes and gene models with DNA repeat, protein similarity and protein domain annotations, using a standard pipeline (see Supplementary Methods for details). Additionally we generate and visualise a series of assembly and gene prediction quality metrics uniformly across the assemblies. For assembly metric visualisation we have introduced data-rich assembly stats plots (described in (Challis in prep)) to provide an overview of a number of key summary statistics in a single graphic (Figure 1).

**Figure1.**
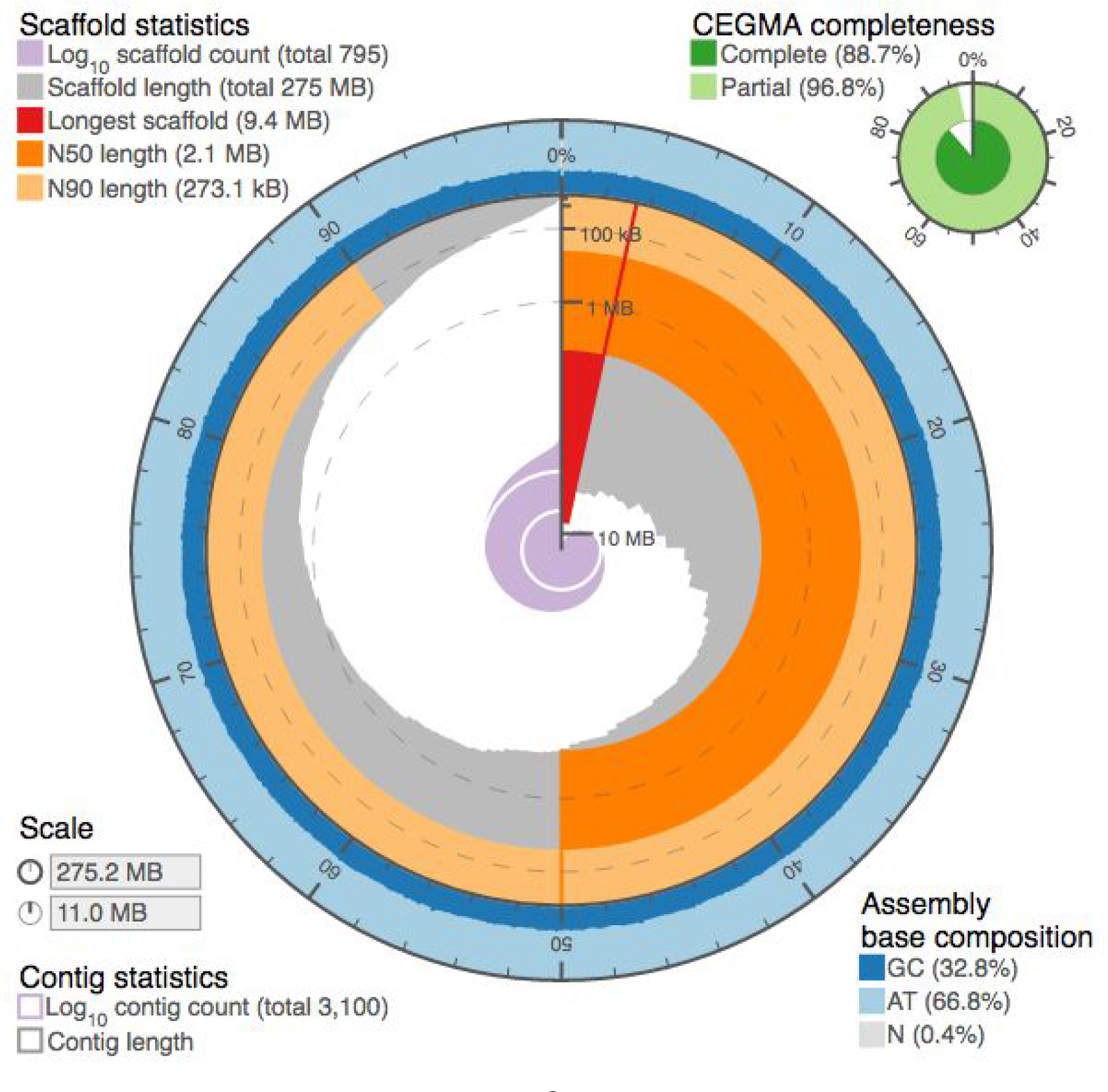
Interactive assembly stats plot (Challis in prep) for Heliconius melpomene assembly Hmel2 (Davey et al. 2016)

As an example of reanalysis of the datasets on Lepbase with consistent methodology, Figure 2 shows a comparison of percent repeat content against genome span for 18 assemblies of varying quality (as indicated by the ratio of N50 to genome size). This shows a positive correlation, as has been observed previously across Metazoa (Canapa et al. 2015), with the largest lepidopteran assembly in the set having > 50% repeat content. Repetitive sequences are modeled and masked using RepeatModeler (Smit AFA, Hubley R 2008–2015) and RepeatMasker (Smit et al. 2013–2015). For each assembly scaffold fasta file a taxon-specific repeat library is generated using RepeatModeler. This library is filtered using the corresponding protein fasta file to remove any hits to annotated proteins that were not annotated as repeats in RepBase. For two species where proteomes are not available for the same assembly we use data from the most closely-related species. Resulting repeat libraries are combined with annotated Lepidoptera repeats from RepBase (Bao et al. 2015) and then used to mask the assembly fasta file. Specific commands used are given in Supplementary Methods. assembly span (Mbp)

**Figure 2.**
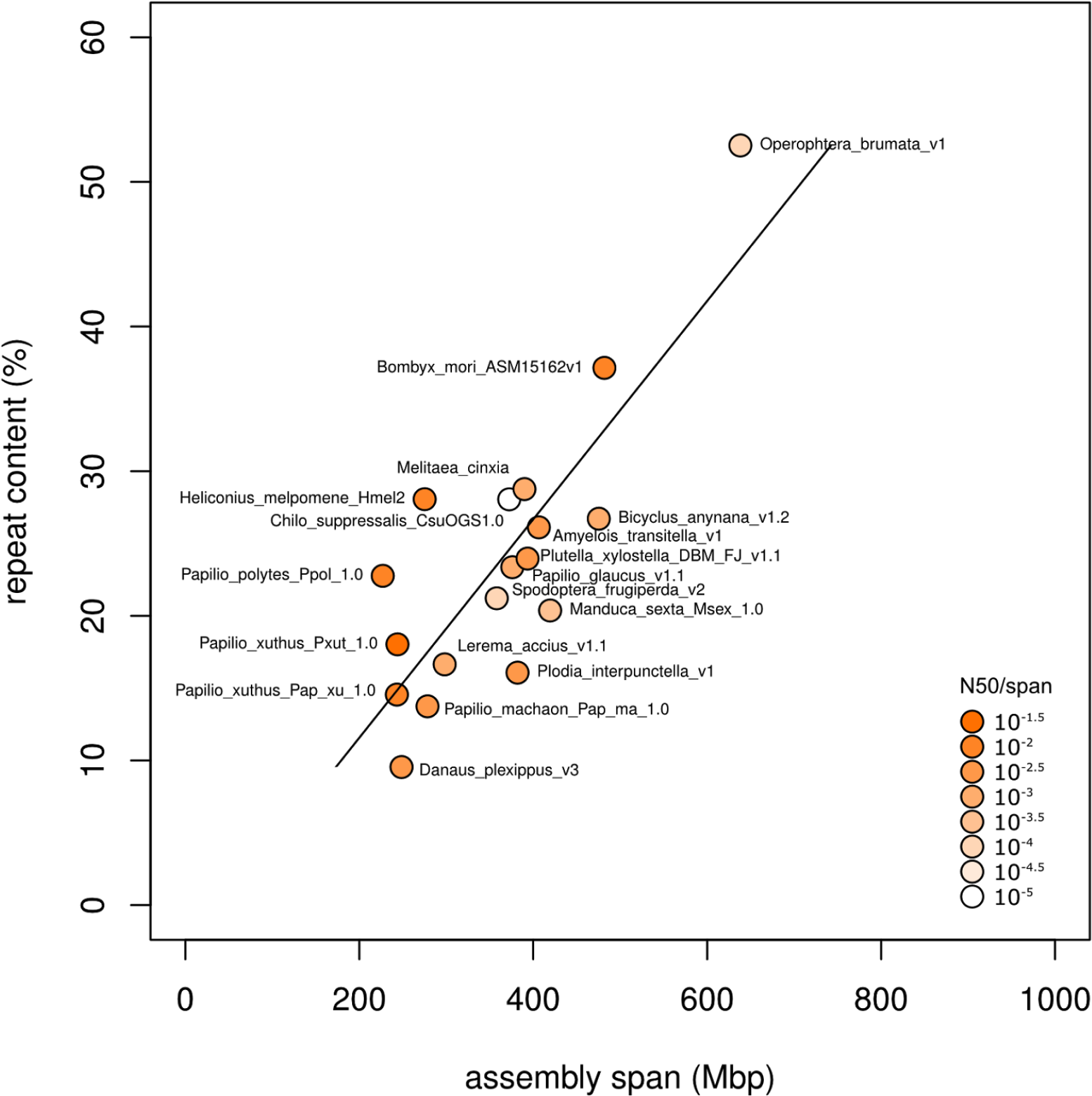
Percent repeat content vs. span for 18 genome assemblies in Lepbase (release v2) showing a positive correlation (y = 0.076⨯ − 3.533, R^2^ = 0.639). Points are shaded according to N50/span as a proxy for assembly quality.

Sequence similarity to proteins in the UniProt/SwissProt database is identified using BLAST+ (Camacho et al. 2009) blastp (the specific commands used are given in Supplementary Methods). Domain annotation of each protein sequence is achieved through the use of InterProScan 5 (Jones et al. 2014) (see Supplementary Methods for code and parameters used). Sequence similarity and domain result files are imported into the Lepbase Ensembl for annotation of genes and visualisation on protein sequences and available for download.

##### Assembly validation using conserved gene sets

CEGMA (Parra et al. 2009) and BUSCO (Simão et al. 2015) both provide an indication of the completeness of an assembly by indicating the proportion of a core set of genes can be identified in an assembly using HMMs. CEGMA has been widely used and offers a single set of 248 core eukaryotic genes (or CEGs) offers a single set of core genes that can be found relatively consistently across eukaryotic assemblies of comparable completeness. We calculate CEGMA scores for each genome, and scores for current Lepbase genomes are presented in Table 1 (full results are available for download at http://download.lepbase.org/v2/cegma). We will offer BUSCO assessment in the future (see below).

#### Infrastructure

A Tier 2 database needs significant compute infrastructure to deliver to a wide community of researchers. Our analyses are undertaken on a dedicated compute cluster (320 cores, up to 1 TB RAM per node) within the Blaxter Lab at the university of Edinburgh. All services are hosted on virtual machines running Ubuntu 14.04 LTS on dedicated servers (12 cores, 48 GB RAM) at the University of Edinburgh or with a cloud hosting provider.

#### Release schedule

As a collection of independent tools and analyses, new data could be added to Lepbase as soon as they become available. However, in order to maintain consistency between tools, it is important to keep the datasets available in each relatively consistent, which is easiest to achieve with a pattern of regular releases, particularly during this initial phase of development when we have been resolving technical challenges while implementing a core toolset. Lepbase v1 was released in October 2015, and v2 followed in February 2016. We plan to continue following a roughly four month release cycle and release v3 in June 2016 - this release will contain 43 genome assemblies in total, alongside major improvements to the comparative analyses that we offer across all assemblies of suitable quality.

#### Impact, Outlook and Future Development

Lepbase is already in use by the lepidopteran research community. Usage statistics (to 20th May 2016) show approximately 260 unique users of the Lepbase Ensembl site since release v1 in October 2015, logging 744 unique sessions with an average session duration of just over 10 minutes. Other components of Lepbase have had 93 (blast.lepbase.org) and 218 (lepbase.org) unique users since release of v2 in February 2016. This usage reflects broad adoption within the relatively small (but growing) lepidopteran research community. The success of Lepbase derives from both our position within the community – from which we are able to work closely with labs to ensure their data are included rapidly – and the maturity and vision of genomics within that the community – including clear requirement for aTier 2 resource.

We have been able to deliver Lepbase effectively by using (and where necessary adapting) existing tools. We have implemented a streamlined Ensembl import pipeline (Challis et al. in prep) that allows us to reduce the time taken to import a new assembly plus annotations. It is thus likely that we will support addition of new data between major releases to respond as quickly as possible to newly available assemblies and encourage rapid data sharing in the Lepidoptera research community.

We are keen to add additional functionality to the database, in particular adding new modes of functional annotation, and also methods for display and exploration of variation data. For example, because CEGMA is no longer being supported, we are collaborating in the development of a lepidopteran-specific BUSCO assembly quality assessment toolkit. BUSCO is a gene-based quality assessment toolkit, similar to CEGMA, that uses a much larger set of genes. BUSCO is more sensitive to the taxonomy of the assembly being assessed, and prior to the release of Lepbase, there were insufficient lepidopteran genomes available for a BUSCO training set to be created. The developers of BUSCO will generate a lepidopteran training set using Lepbase, and we will analyse all our assemblies using BUSCO in future. With these and other tools we will continue to serve the current research community and attract new researchers to the rich genomic resources for Lepidoptera.

## Availability and resources

All lepbase web sites and services are available for use without restriction, apart from webapollo.lepbase.org for which registration is required before editing gene models. All source code written for this project is available in open source repositories under MIT or Apache 2.0 licenses: **lepbase.org** uses a custom wordpress theme, available at https://github.com/lepbase/lepbase-theme: **ensembl.lepbase.org** uses custom Ensembl plugins, available at https://github.com/lepbase/lepbase-ensembl, https://github.com/lepbase/lepbase-search, and https://github.com/lepbase/lepbase-blast-linkout: **blast.lepbase.org** uses a fork of the SequenceServer source code (https://github.com/wurmlab/sequenceserver), available at https://github.com/lepbase/lepbase-sequenceserver.

## Availability of supporting data

All results and datafiles are available at http://download.lepase.org

## Acknowledgements

Lepbase is funded by a Biotechnology and Biological Sciences Research Council Tools and Resources fund award (<u>BB/K020161/1</u>, <u>BB/K019945/1</u>, <u>BB/K020129/1</u>) to MB, CDJ and KD. We wish to thank members of the Lepidopteran research community for providing the data on Lepbase, in particular those who have allowed us to include unpublished data: Paul Brakefield, Ben Elsworth, Jim Mallet, Reuben Nowell, Steve Paterson and Hugh Robertson. We also wish to thank members of the Ensembl, VectorBase and WormBase Parasite teams for helpful discussions.

